# CLASP2 lattice-binding near microtubule plus ends stabilizes kinetochore attachments

**DOI:** 10.1101/634907

**Authors:** Hugo Girão, Naoyuki Okada, Ana C. Figueiredo, Zaira Garcia, Tatiana Moutinho-Santos, Jorge Azevedo, Sandra Macedo-Ribeiro, Ikuko Hayashi, Helder Maiato

## Abstract

The fine regulation of kinetochore microtubule dynamics during mitosis ensures proper chromosome segregation by promoting error correction and spindle assembly checkpoint (SAC) satisfaction. CLASPs are widely conserved microtubule plus-end-tracking proteins that regulate microtubule dynamics throughout the cell cycle and independently localize to kinetochores during mitosis. Thus, CLASPs are ideally positioned to regulate kinetochore microtubule dynamics, but the underlying molecular mechanism remains unknown. Here we found that human CLASP2 can dimerize through its C-terminal kinetochore-targeting domain, but kinetochore localization was independent of dimerization. CLASP2 kinetochore localization, microtubule plus-end-tracking and microtubule lattice binding through TOG2 and TOG3 (but not TOG1) domains, independently sustained normal spindle length, timely SAC satisfaction, chromosome congression and faithful segregation. Measurements of kinetochore microtubule half-life in living cells expressing RNAi-resistant mutants revealed that CLASP2 kinetochore localization, microtubule plus-end-tracking and lattice binding cooperatively modulate kinetochore microtubule stability during mitosis. Thus, CLASP2 regulates kinetochore microtubule dynamics by integrating distinctive microtubule-binding properties at the kinetochore-microtubule interface to ensure chromosome segregation fidelity.

## Introduction

Microtubule dynamics are modulated by several Microtubule-Associated Proteins (MAPs) (Maiato et al., 2004). Some MAPs specifically accumulate at the growing plus ends of microtubules and are collectively known as microtubule plus-end-tracking proteins or +TIPs (Akhmanova and Steinmetz, 2008). CLIP-associated proteins (CLASPs) are widely conserved +TIPs that stabilize microtubule plus ends by suppressing catastrophes and promoting rescue (Aher et al., 2018; Al-Bassam et al., 2010; Lawrence et al., 2018; Majumdar et al., 2018). Humans have two CLASP paralogues, CLASP1 and CLASP2, which exist as multiple isoforms (Akhmanova et al., 2001; Inoue et al., 2000; Lemos et al., 2000). Structurally, CLASPs harbor three distinct functional domains: 1) a characteristic basic region, also found in other +TIPs, comprising two serine-x-isoleucine-proline (SxIP) motifs that enable microtubule plus-end-tracking via interaction with EB proteins (Honnappa et al., 2009; Mimori-Kiyosue et al., 2005); 2) the presence of two to three tumor overexpression gene (TOG) domains, which interact with the microtubule lattice by distinguishing the structural conformation of the αβ-tubulin dimer on microtubule protofilaments (Leano et al., 2013; Maki et al., 2015); and 3) a C-terminal domain required for kinetochore localization (Maia et al., 2012; Maiato et al., 2003a; Mimori‐Kiyosue et al., 2006) and protein dimerization (Al-Bassam et al., 2010; Funk et al., 2014; Patel et al., 2012), as well as interaction with other kinetochore proteins, including CLIP170, CENP-E and Plk1 (Akhmanova et al., 2001; Dujardin et al., 1998; Maffini et al., 2009; Maia et al., 2012).

During mitosis, mammalian CLASPs play redundant roles in the organization of the mitotic spindle, and interference with their function results in several mitotic abnormalities, including monopolar, short and multipolar spindles (Logarinho et al., 2012; Maiato et al., 2003a; Maiato et al., 2003b; Mimori‐Kiyosue et al., 2006; Pereira et al., 2006). CLASPs localize at the fibrous corona region of the kinetochore throughout mitosis, where they play a critical role in the regulation of kinetochore microtubule dynamics required for microtubule poleward flux and turnover, as well as the correct alignment and segregation of chromosomes (Logarinho et al., 2012; Maffini et al., 2009; Maiato et al., 2003a; Maiato et al., 2005; Maiato et al., 2003b; Pereira et al., 2006). Previous works revealed that, during prometaphase, CLASP1 interacts with the kinesin-13 Kif2B to promote kinetochore microtubule turnover, which is necessary for the correction of erroneous attachments (Maffini et al., 2009; Manning et al., 2010). In metaphase, CLASP1 interacts with Astrin, which promotes kinetochore microtubule stabilization required for spindle assembly checkpoint (SAC) satisfaction (Manning et al., 2010). This places CLASP1 as part of a regulatory switch that enables the transition between labile-to-stable kinetochore microtubule attachments by establishing temporally distinct interactions with different partners at the kinetochore. CLASP2 is regulated by Cdk1 and Plk1 phosphorylation as cells progressively reach metaphase, leading to a gradual stabilization of kinetochore-microtubule attachments (Maia et al., 2012). While this broad picture provides important information on the molecular context in which CLASPs operate at the kinetochore-microtubule interface, we still lack a detailed mechanistic view on how the intrinsic properties of CLASPs modulate kinetochore microtubule dynamics.

Here we focused on human CLASP2 to investigate how its distinct functional domains affect mitosis, with emphasis on the regulation of kinetochore microtubule dynamics. Our findings revealed that CLASP2 integrates multiple independent features, including microtubule plus-end-tracking, microtubule lattice binding through TOG domains and kinetochore localization, to modulate kinetochore microtubule dynamics required for faithful chromosome segregation during mitosis in human cells.

## Results and Discussion

### CLASP2 can form dimers through its C-terminal domain

Previous studies have come to contradictory conclusions regarding the native structure of CLASPs. Based on the hydrodynamic profile of several kinetochore sub-complexes present in *Xenopus* egg extracts, it was proposed that the single *Xenopus* CLASP exists as an elongated monomer in solution (Emanuele et al., 2005). This was subsequently confirmed by Fluorescence Correlation Spectroscopy and Photon Counting for human GFP-CLASP2 (Drabek et al., 2006), as well as single molecule fluorescence intensity quantification of recombinant human GFP-CLASP2 (Aher et al., 2018). However, another set of studies with recombinant human CLASP1 and its counterparts in *S. pombe* and *S. cerevisiae* indicated that they can form homodimers in solution through their C-terminal domain (Al-Bassam et al., 2010; Funk et al., 2014; Patel et al., 2012). To shed light into this debate, we investigated the native state of CLASP2 in human cells. This was initially assessed by blue native polyacrylamide gel electrophoresis followed by western-blot analysis (Fig. 1A). We observed a fast migrating band, consistent with a monomeric form of CLASP2, as well as higher molecular weight species that may have arisen from formation of homodimers, heterodimers with CLASP1 and higher order multiprotein complexes. Interestingly, the migration profile of CLASP2 was similar in mitotic and asynchronous cell populations, indicating that the previously reported CLASP2 phosphorylation in mitosis (Maia et al., 2012) does not significantly alter any potential oligomerization or higher order state of CLASP2.

**Figure 1.**
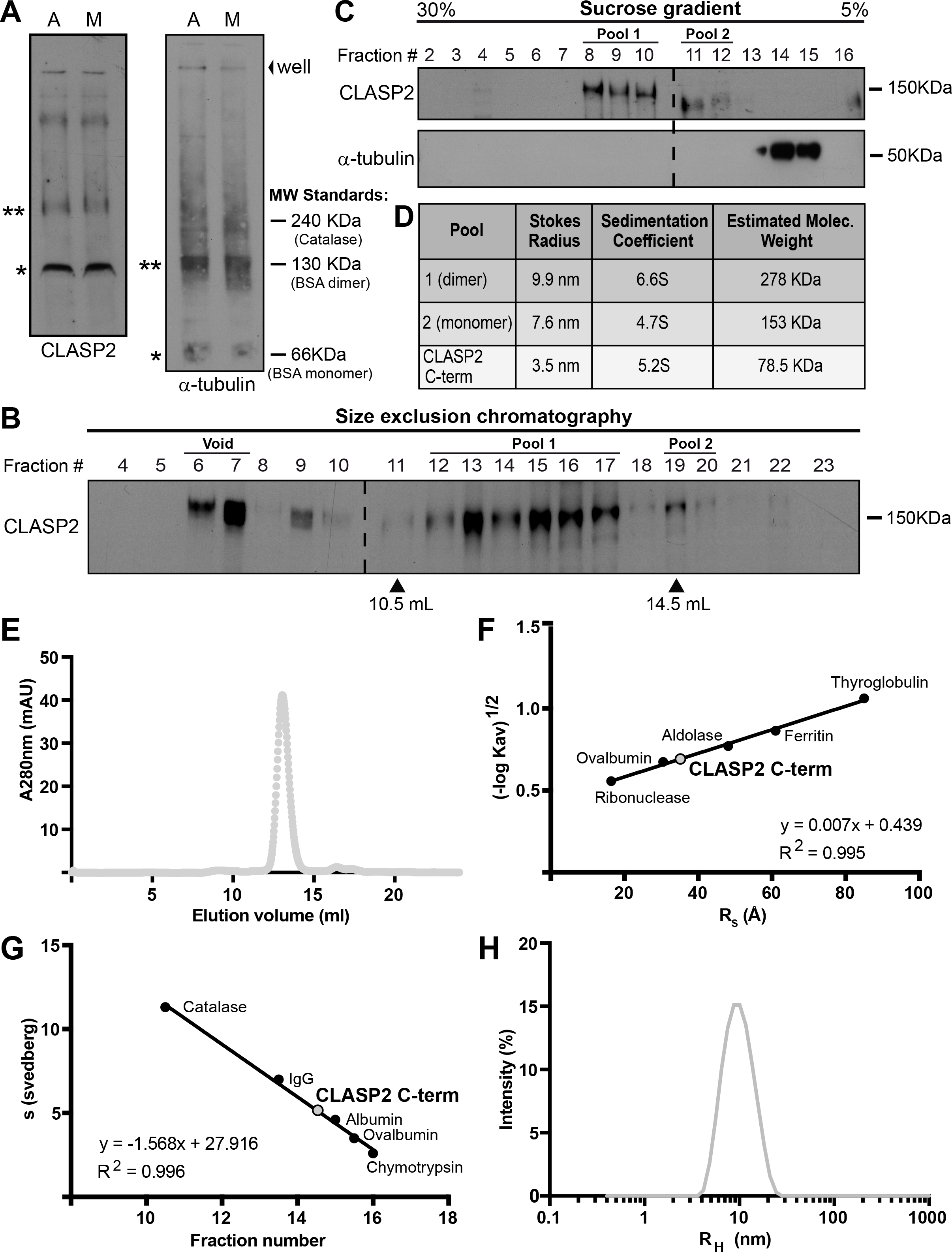
CLASP2 can form dimers through its C-terminal kinetochore-targeting domain. **(A)** Extracts from asynchronous (A) and mitotic (M) HeLa cells resolved by BN-PAGE and analyzed by western blot for CLASP2 (left) or α-tubulin (right). Indicated apparent molecular weights were estimated by comparison with the gel mobility behavior of protein standards of known molecular weights, as indicated. Dimerization of α-tubulin (130 kDa) was used as a control. Single or double asterisks indicate the position of putative monomers and dimers, respectively. **(B)** Mitotic enriched HeLa cell extracts were separated by size exclusion chromatography. Collected fractions were TCA-precipitated and analyzed by western blot with anti-CLASP2 antibodies. **(C)** Mitotic enriched HeLa cell extracts were loaded on a 5%-30% sucrose gradient. Fractions collected after centrifugation were TCA-precipitated and analyzed by western blot with the indicated antibodies. Note that CLASP2 and free tubulin are collected in distinct fractions. **(D)** Summary of Stokes radius (nm), sedimentation coefficients (S) and estimated molecular weights (KDa) calculated for the different CLASP2 forms**. (E)** Size exclusion chromatogram of human recombinant CLASP2 C-terminal, eluting in a single peak. **(F)** Size exclusion chromatography calibration curve used to determine the Stoke’s radius of CLASP2 C-terminal. **(G)** Density gradient centrifugation curve used to determine the sedimentation coefficient of recombinant CLASP-2 C-terminal. **(H)** Dynamic light scattering analysis of the CLASP2 C-terminal showing an hydrodynamic radius of 36.1 Å, corresponding to an estimated molecular weight of 67.9 KDa.

Next, we calculated the molecular weight of CLASP2 from mitotic cells using the method of Siegel and Monty (Siegel and Monty, 1966), which combines the Stokes’ radius obtained from size exclusion chromatography (Fig. 1B) and the sedimentation coefficients derived from density gradient centrifugation (Fig. 1C). Accordingly, we identified two pools of CLASP2 with apparent molecular weights of 153 kDa and 278 kDa, respectively (Fig. 1D), which did not co-elute with α-tubulin (Fig. 1C), suggesting that, unlike its yeast counterparts (Al-Bassam et al., 2010; Funk et al., 2014; Majumdar et al., 2018), human CLASP2 is not normally associated with free tubulin, in agreement with recent in vitro reconstitution experiments (Aher et al., 2018; Maki et al., 2015). These results are consistent with the existence of monomeric (theoretical value for the longest CLASP2α isoform = 166 kDa) and dimeric forms of CLASP2 in human cells. Further calculation of the frictional ratio of these two pools yielded a value of ~2, indicating that CLASP2 molecules are non-globular and considerably elongated.

To determine whether CLASP2 dimerization is mediated by the C-terminal region, as previously reported for human CLASP1 and its counterparts in yeasts (Al-Bassam et al., 2010; Funk et al., 2014; Patel et al., 2012), we investigated the oligomerization of a C-terminal recombinant fragment of human CLASP2 (CLASP2-1213-1515). CLASP2 C-terminal eluted as a single peak after size exclusion chromatography (Fig. 1E), in a volume corresponding to a Stokes’ radius of 35.1Å (Fig. 1F). Density gradient centrifugation analysis revealed a sedimentation coefficient of 5.2 (Fig. 1G). Using the Siegel and Monty method, the calculated molecular weight was 78.5 kDa, consistent with dimer formation (theoretical value 75.5 kDa) (Fig. 1D). Analysis of CLASP2 C-terminal by Dynamic Light Scattering (DLS), which measures Brownian motion and relates this to the size of particles in solution, also revealed a single population with low polydispersity values (10%) and a hydrodynamic radius of 36.1Å (Fig. 1H), corresponding to an estimated molecular weight of 67.9 kDa. Taken together, these results indicate that human CLASP2 can form dimers through its C-terminal domain. The existence as monomer or dimer in solution or in cells might depend on the physiological concentration of CLASP2 and its accumulation in particular cellular structures (e.g. kinetochores).

### CLASP2 C-terminal domain is required for kinetochore localization independently of dimerization

To investigate how CLASP2 impacts kinetochore microtubule dynamics during mitosis, we used an RNAi-based rescue system with RNAi-resistant mRFP-CLASP2γ-derived constructs harboring specific mutations that disrupt the different CLASP2 functional domains (Maki et al., 2015), either alone or in combination (Fig. 2A). In addition to the wild type (WT) mRFP-CLASP2γ construct, we used a construct mutated in both SxIP motifs (IP12) (Honnappa et al., 2009) that disrupts CLASP2-EB protein interaction and consequently microtubule plus-end-tracking; a construct mutated in both TOG2 and TOG3 domains (2ea-3eeaa) (Al-Bassam et al., 2007; Leano et al., 2013) required for microtubule lattice binding; as well as a construct combining mutations at both SxIP motifs and in TOG3 (IP12-3eeaa). Additionally, we used a construct with a truncated C-terminal domain (ΔC), which mediates CLASP2 dimerization (this study) and kinetochore localization (Maia et al., 2012; Mimori‐Kiyosue et al., 2006). Finally, to dissect the requirement of CLASP2 dimerization for its kinetochore localization, we generated a construct in which the human CLASP2 C-terminal was replaced by an artificial dimerization domain based on the Gcn4-Leucine Zipper sequence (ΔC-Gcn4), as shown previously for the *S. cerevisiae* CLASP orthologue Stu1 (Funk et al., 2014).

**Figure 2.**
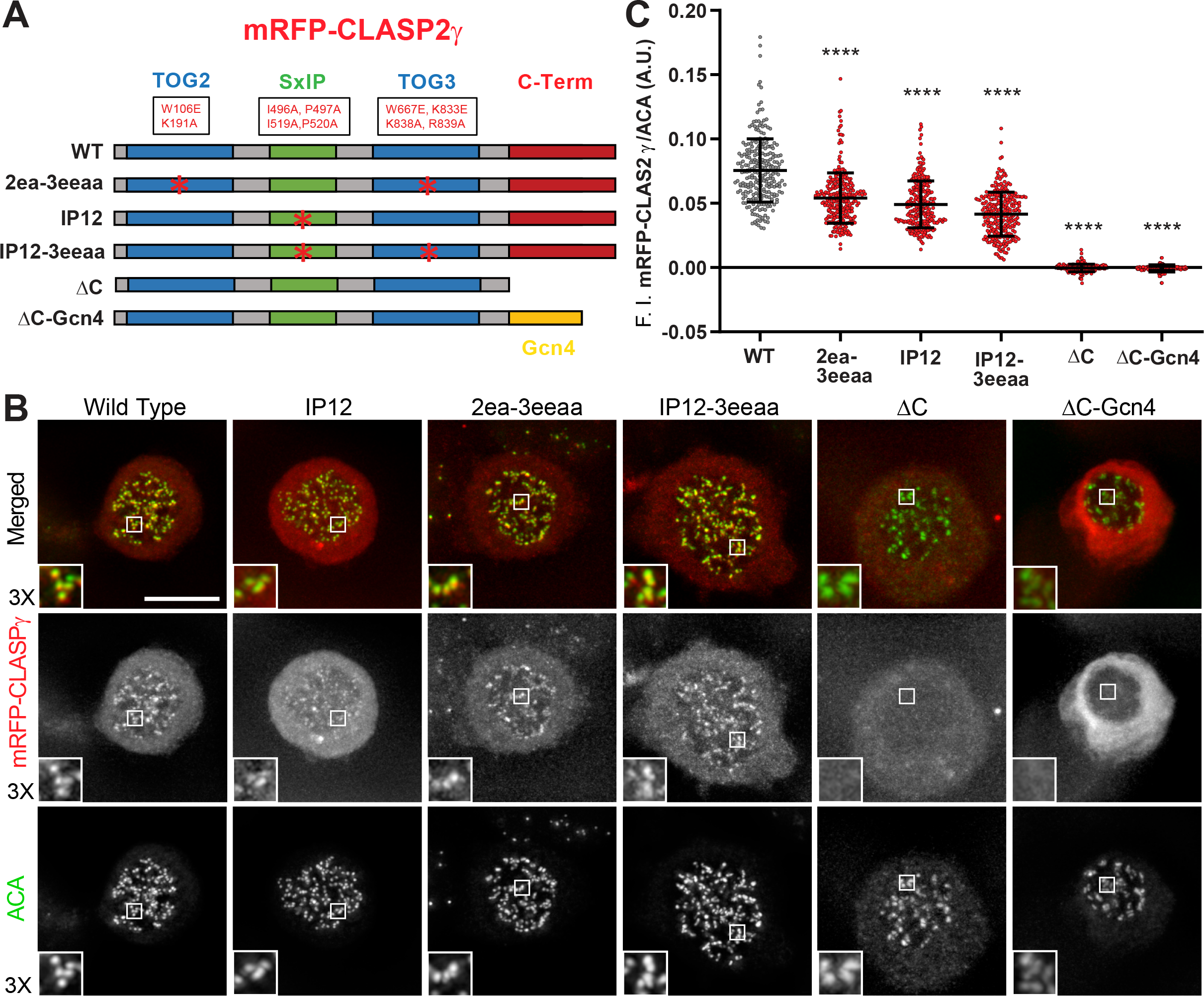
CLASP2 C-terminal domain is required for kinetochore localization independently of dimerization. **(A)** Schematic representation of the RNAi-resistant mRFP-CLASP2γ constructs used in this project: Wild Type (WT) CLASP2γ; the mutant constructs IP12, 2ea-3eeaa, IP12-3eeaa; the truncated version at C-terminal ΔC; and the hybrid construct ΔC-Gcn4. Mutations in TOG2 and TOG3 domains and SxIP motifs are indicated. **(B)** Immunofluorescence analysis of the U2OS PA-GFP-α-tubulin and mRFP-CLASP2γ transduced cell lines, arrested in mitosis by the addition of nocodazole to depolymerize microtubules, showing the localization of the different CLASP2 constructs at kinetochores, except the ΔC and ΔC-Gcn4 constructs. Cells were immunostained with anti-centromere antibodies (ACA) from CREST patients (green) and the mRFP signal of the different CLASP2 constructs (red). Detailed images of mRFP and ACA co-localization are magnified 3X. Scale bar is 5 μm. **(C)** Quantification of kinetochore fluorescence intensity (F.I.) of mRFP/ACA signals for each cell line after background subtraction. Each data point represents an individual kinetochore. Bars represent mean and standard deviation. Quantifications from 3 independent experiments: WT = 26 cells, 260 kinetochores; 2ea-3eeaa = 27 cells, 267 kinetochores; IP12 = 27 cells, 269 kinetochores, IP123eeaa = 26 cells, 253 kinetochores ΔC = 26 cells, 253 kinetochores ΔC-Gcn4 = 18 cells, 196 kinetochores. All mutant constructs showed a decreased intensity of the signal at the kinetochores compared with WT. **** = p<<0.0001.

After stable transduction using a lentiviral system in a human U2OS cell line stably expressing photoactivatable (PA) GFP-α-tubulin (Ganem et al., 2005), all constructs expressed at comparable levels (Sup. Fig. 1). We then determined the localization of the different constructs after RNAi-mediated depletion of both CLASP1 and CLASP2 (Sup. Fig. 2) to avoid functional redundancy (Mimori‐Kiyosue et al., 2006; Pereira et al., 2006) and potential dimerization with the endogenous proteins. As expected, the IP12 construct failed to localize at microtubule plus ends, as determined by co-localization with EB1 in interphase cells (Sup. Fig. 3A), whereas the 2ea-3eeaa construct failed to associate with mitotic spindle microtubules (Sup. Fig. 3B). In agreement, the IP12-3eeaa construct showed a compromised localization at both microtubule plus ends and spindle microtubules (Sup. Fig. 3A, B). To investigate the localization of the different CLASP2 constructs at unattached kinetochores we used mitotic cells treated with nocodazole to depolymerize spindle microtubules. All constructs, except the ΔC and ΔC-Gcn4, were able to localize at unattached kinetochores (Fig. 2B). In order to substantiate these findings, we quantified the fluorescence intensity of each CLASP2 construct relative to the signal from constitutive anti-centromere antibodies from CREST patients. As expected, mutant CLASP2-ΔC and CLASP2-ΔC-Gcn4 proteins were virtually undetectable at unattached kinetochores (Fig. 2C), confirming that the C-terminal domain of CLASP2 is necessary for kinetochore localization, while demonstrating that dimerization *per se* is not sufficient to target CLASP2 to kinetochores. This reveals a critical difference between human CLASP2 and its orthologue Stu1 in *S. cerevisiae*, in which dimerization was shown to be sufficient for kinetochore targeting (Funk et al., 2014). Surprisingly, although all other CLASP2 mutants that compromise microtubule plus-end-tracking and association with the microtubule lattice were detectable at unattached kinetochores, they all showed a statistically significant decrease (~20%) relative to WT CLASP2 (Fig. 2C). We attribute this reduction to slight differences in the expression of the distinct constructs and/or possible alterations in the tridimensional conformation/folding of CLASP2 that might compromise its normal interaction with kinetochores.

### Regulation of mitotic spindle length by CLASP2 depends on its kinetochore localization and microtubule-binding properties

CLASPs have been previously implicated in the control of mitotic spindle length through their role in the incorporation of tubulin required for kinetochore microtubule poleward flux (Logarinho et al., 2012; Maffini et al., 2009; Maiato et al., 2003a; Maiato et al., 2005; Maiato et al., 2002; Mimori‐Kiyosue et al., 2006). However, it remains unknown how CLASPs mediate this process. One possibility is that, similar to other TOG-domain containing proteins that localize to kinetochores (Ayaz et al., 2012; Geyer et al., 2018; Miller et al., 2016; Nithianantham et al., 2018), CLASPs act as microtubule polymerases by interacting directly with free tubulin in solution. Another possibility is that CLASPs regulate spindle microtubule flux by controlling the interaction between kinetochores and the microtubule lattice, near polymerizing microtubule plus-ends. To distinguish between these possibilities, we used fluorescence microscopy in fixed cells and the same RNAi-rescue strategy to investigate how the different CLASP2 functional domains contribute to ensure normal mitotic spindle length. As expected, RNAi-mediated depletion of endogenous CLASPs from U2OS cells resulted in shorter spindles (Maffini et al., 2009), and expression of RNAi-resistant WT CLASP2γ completely rescued normal spindle length (Fig 3A, C). This further demonstrates CLASPs functional redundancy, while revealing that the TOG1 domain that is lacking in the CLASP2γ isoform is completely dispensable for the regulation of mitotic spindle length. In contrast, expression of all the other mutant constructs failed to rescue normal spindle length (Fig. 3A, C). Interestingly, disruption of the TOG2 and TOG3 domains affected spindle length to a similar extent as disruption of the SxIP motifs, whereas the combined disruption of both TOG3 domain and SxIP motifs exacerbated this effect almost comparably with the disruption of the kinetochore-targeting domain of CLASP2 or CLASPs depletion alone (Fig. 3A, C). Altogether, these results indicate that microtubule plus-end-tracking and microtubule lattice-binding through the TOG2 and TOG3 domains are independently required to ensure normal spindle length. However, these properties per se are insufficient to ensure the normal function of CLASP2, which requires targeting to kinetochores, independently of dimerization. Finally, expression of all mutant CLASP2 constructs resulted in a significant increase in anaphase and telophase cells with chromosome missegregation events (Fig. 3B, D), suggesting a role for CLASP2 in error correction.

**Figure 3.**
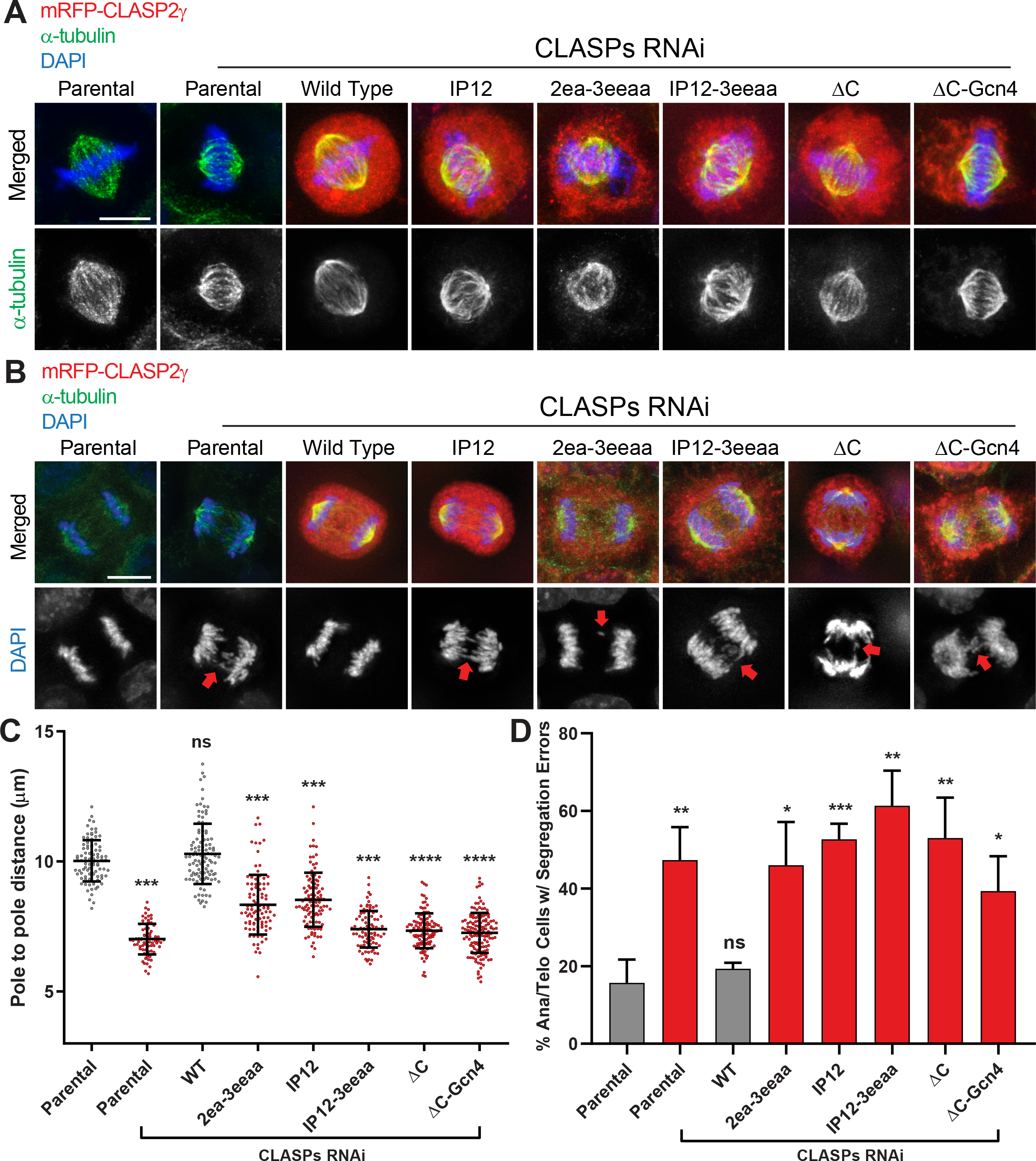
Regulation of mitotic spindle length by CLASP2 depends on its kinetochore localization and microtubule-binding properties. **(A)** Immunofluorescence analysis of the U2OS PA-GFP-α-tubulin cell lines in metaphase. Parental (without exogenous mRFF-CLASP2γ constructs) and mutant cell lines depleted from endogenous CLASPs with siRNA #12A, showed shorter spindles relative to cells expressing WT mRFF-CLASP2γ. Scale bar is 5 μm. **(B)** Immunofluorescence analysis of the U2OS PA-GFP-α-tubulin cell lines in anaphase. Segregation errors are indicated with red arrows in the DAPI channel. Parental and CLASP2 mutant cell lines depleted of endogenous CLASPs by siRNA #12A showed increased chromosome segregation errors relative to WT. Scale bar is 5 μm. **(C)** Quantification of mitotic spindle length in each cell line: the first column represents parental U2OS PA-GFP-α-tubulin cell line without RNAi treatment, the remaining columns represent treatment with siRNA #12A. Each data point represents an individual cell. Bars represent mean and standard deviation. Quantifications from 3 independent experiments: Parental without RNAi = 88 cells; Parental with RNAi = 74 cells; WT = 108 cells; 2ea-3eeaa = 96 cells; IP12 = 105 cells; IP12-3eeaa = 88 cells; ΔC = 112 cells; ΔC-Gcn4 = 142 cells. Parental and mutant cell lines depleted of endogenous CLASPs show a statistical significant decrease in spindle length. ns = non-significant, *** = p<0.001, **** = p<0.0001. **(D)** Quantification of chromosome segregation errors in anaphase and telophase cells: each column represents the percentage of anaphase and telophase cells with at least one segregation error; the first column represents parental U2OS PA-GFP-α-tubulin cell line without RNAi treatment, the remaining columns represent treatment with the siRNA #12A. Quantifications from 3 independent experiments: Parental without RNAi = 92 cells; Parental with RNAi = 117 cells; WT = 153 cells; 2ea-3eeaa = 126 cells; IP12 = 135 cells; IP12-3eeaa = 121 cells; ΔC = 120 cells; ΔC-Gcn4= 106 cells. CLASPs-depleted cells and mutant cell lines showed higher percentage of cells with segregation errors. ns = non-significant, * = p<0.05; ** = p< 0.01; *** = p<0.001.

### Chromosome congression, timely SAC satisfaction and chromosome segregation fidelity rely on both CLASP2 kinetochore- and microtubule-binding properties

To gain a deeper insight on how the distinct CLASP2 functional domains impact mitosis we resorted to live-cell imaging analysis by spinning-disc confocal microscopy. To monitor chromosome movements we transduced H2B-GFP in each of the stable U2OS lines expressing the distinct CLASP2 constructs and added SiR-tubulin at 20 nM to visualize microtubules (Lukinavicius et al., 2014). As expected, CLASPs depletion from U2OS cells resulted in a significant delay from nuclear envelope breakdown (NEB) to anaphase onset relative to controls (Fig 4A, B), and this delay was fully rescued by expression of the WT CLASP2γ construct, but not by any of the CLASP2 mutant constructs (Fig 4A, B). These results suggest that CLASP2 at the kinetochore requires microtubule plus-end-tracking and lattice binding capacities to establish a functional kinetochore-microtubule interface necessary for timely SAC satisfaction.

**Figure 4.**
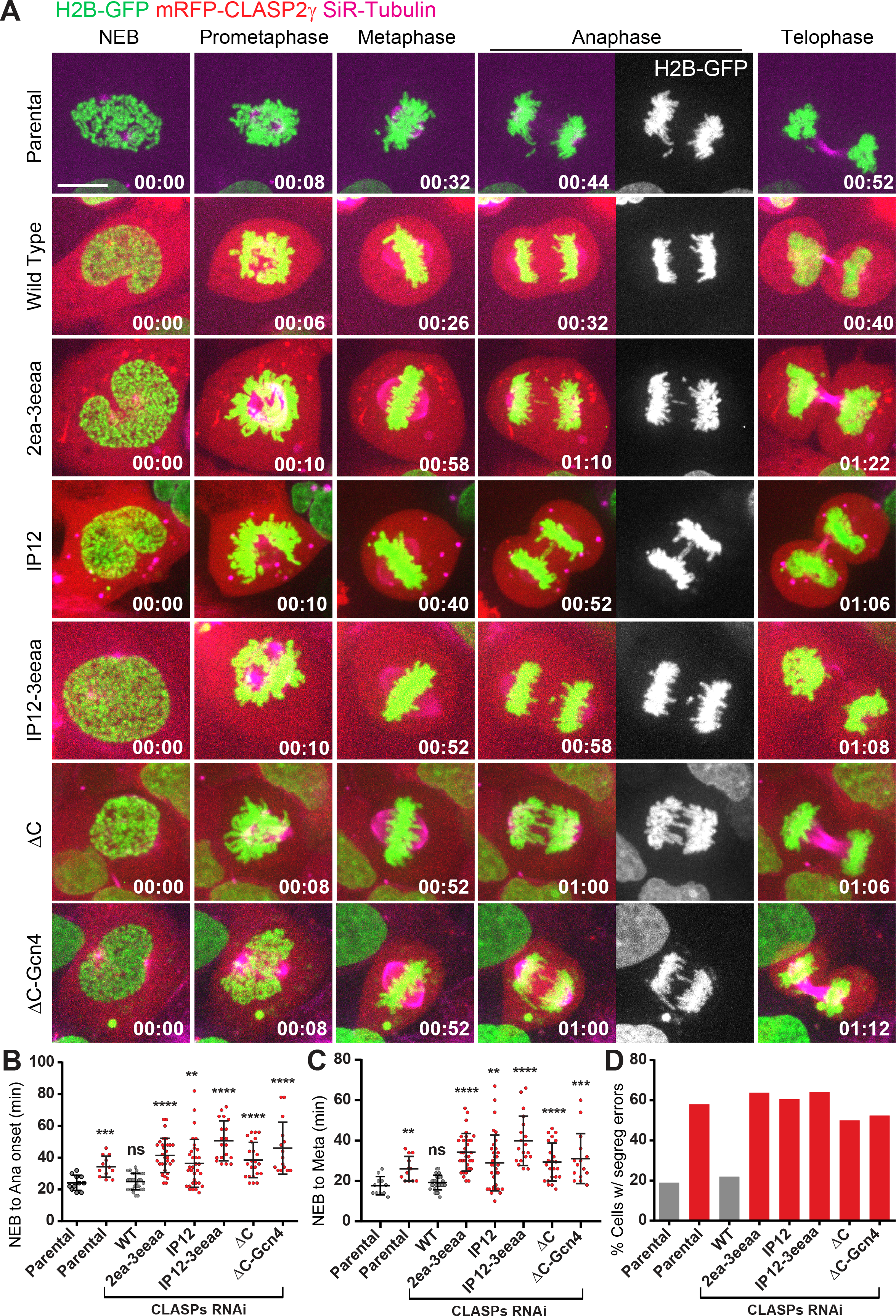
Chromosome congression, timely SAC satisfaction and chromosome segregation fidelity relies on both CLASP2 kinetochore- and microtubule-binding properties. **(A)** Live-cell imaging of the parental U2OS PA-GFP-α-tubulin cells and respective cell lines expressing the mRFP-CLASP2γ constructs, after endogenous depletion of CLASPs with RNAi. mRFP-CLASPγ (red), DNA (blue) and SiR-tubulin (magenta) are depicted. Timing 00:00 starts at Nuclear Envelope Breakdown (NEB). The anaphase panel additionally shows the H2B-GFP signal to highlight the segregation errors. Time is in hours:minutes. Scale bar is 5 μm. **(B)** Mitotic timing quantification between NEB and Anaphase (Ana) onset. First column represents parental U2OS PA-GFP-α-tubulin cells. The remaining columns represent CLASPs RNAi after treatment with siRNA #12A. Each data point represents an individual cell. Bars represent mean and standard deviation. Parental without RNAi = 11 cells, 3 independent experiments; Parental with RNAi = 12 cells, 3 independent experiments; WT = 28 cells, 6 independent experiments; 2ea-3eeaa = 32 cells, 14 independent experiments; IP12 = 33 cells, 13 independent experiments; IP12-3eeaa = 19 cells, 7 independent experiments; ΔC = 23 cells, 9 independent experiments; ΔC-Gcn4 = 15 cells, 5 independent experiments. Parental and mutant cells depleted of endogenous CLASPs showed a significant increase in mitotic duration. ns = non-significant, ** = p<0.01, *** = p<0.001, **** = p<0.0001. **(C)** Quantification of the time between NEB and Metaphase (Meta) for the same data set as in B. First column represents parental U2OS PA-GFP-α-tubulin cells. The remaining columns represent CLASPs RNAi after treatment with siRNA #12A. A significant delay in chromosome congression is observed in parental and mutant cell lines after CLASPs depletion. ns = non-significant, ** = p<0.01, *** = p<0.001, **** = p<0.0001. **(D)** Quantification of segregation errors during live cell imaging for the same data set as in B and C; each column represents the percentage of cells with at least one segregation error detected in the total cells imaged for each cell line. Parental and mutant cells depleted of endogenous CLASPs showed a higher frequency of cells with segregation errors.

Measurement of the duration from NEB to metaphase in the same experimental setup revealed that the observed delay in anaphase onset after CLASPs depletion, or inefficient rescue with the different CLASP2 mutant constructs, was essentially due to a delay in completing chromosome congression to the spindle equator (Fig. 4A, C), consistent with a role of CLASP2 in the regulation of kinetochore-microtubule attachments. Moreover, this delay correlated with an increase in chromosome segregation errors during anaphase and telophase (Fig. 4D), in agreement with our fixed cell analysis (Fig. 3D). Taken together, these results strongly suggest that multiple domains of CLASP2 ensure normal kinetochore microtubule dynamics required for chromosome congression, timely SAC satisfaction and efficient error correction during mitosis.

### Multimodal regulation of kinetochore-microtubule dynamics by CLASP2

To directly test how the different CLASP2 domains impact kinetochore microtubule dynamics we quantified kinetochore and non-kinetochore microtubule half-life by Fluorescence Dissipation after Photoactivation (FDAPA) (Ferreira et al., 2018). By fitting the fluorescence decay over time to a double exponential curve, it is possible to discriminate two spindle microtubule populations with fast and slow turnover that are thought to correspond to the less stable non-kinetochore microtubules and more stable kinetochore microtubules, respectively (Bakhoum et al., 2009a; Bakhoum et al., 2009b; Zhai et al., 1995). Admittedly, while this approach is unable to distinguish multiple stable microtubule populations that might exist in the spindle, it remains the golden standard to detect subtle alterations in kinetochore microtubule dynamics that cannot be disclosed by less sensitive methods, such as cold-shock or treatment with microtubule poisons. As so, depletion of endogenous CLASPs by RNAi resulted in a significant decrease in kinetochore microtubule half-life during metaphase when compared to controls (Fig. 5A, B), without significantly affecting the half-life of non-kinetochore microtubules (Fig. 5A, C). Interestingly, previous measurements of kinetochore microtubule half-life after CLASPs depletion from human U2OS cells suggested a role in kinetochore microtubule destabilization (Maffini et al., 2009). However, this previous study did not distinguish between late prometaphase and metaphase cells, reinforcing the idea that CLASPs are part of a regulatory switch that controls the transition between labile-to-stable kinetochore microtubule attachments (Maia et al., 2012; Manning et al., 2010). Importantly, the WT CLASP2γ construct was able to completely rescue normal kinetochore microtubule half-life (Fig. 5A, B), indicating full functionality and dispensability of the TOG1 domain for the stabilization of kinetochore microtubules during metaphase. In contrast, mutation of the TOG2 and TOG3 domains, or the SxIP motifs of CLASP2, resulted in less stable kinetochore microtubules and consequently incapacity to rescue the effect caused by CLASPs depletion, a tendency that was exacerbated by the combined mutation of the TOG3 and SxIP domains (Fig. 5A, B). These results suggest that the microtubule lattice binding properties of CLASP2 act synergistically with its microtubule plus-end-tracking ability to stabilize kinetochore-microtubule attachments during metaphase in human cells.

**Figure 5.**
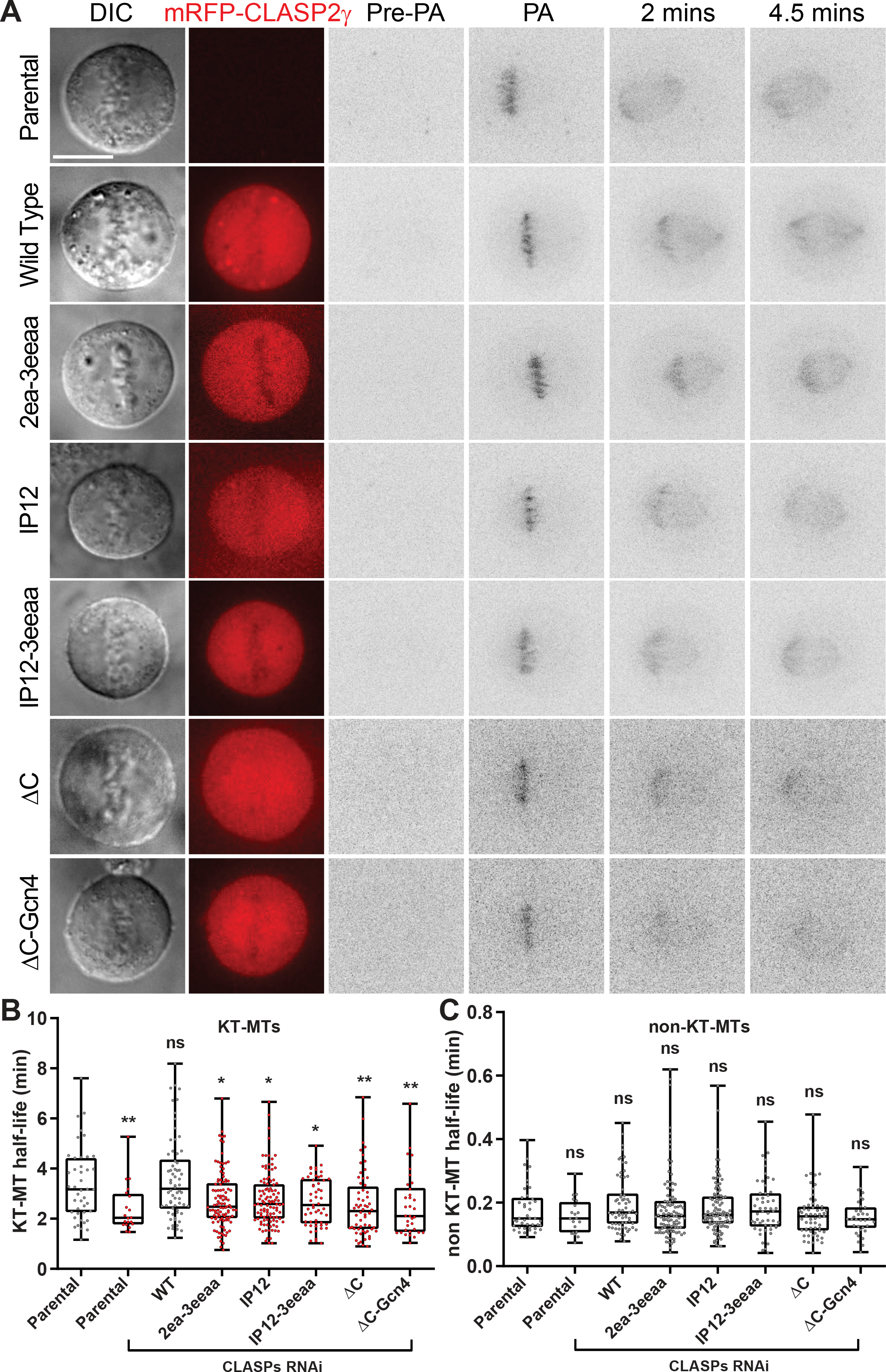
Multimodal regulation of kinetochore microtubule dynamics by CLASP2. **(A)** Photoactivation experiments in live U2OS parental PA-GFP-α-tubulin cells and expressing the different mRFP-CLASP2γ constructs, after RNAi against endogenous CLASPs; the expression of PA-GFP-α-tubulin enables the activation of a stripe near the metaphase plate by a near-UV laser. Panels represent DIC, the mRFP-CLASP2γ signal (red) and the PA-GFP-α-tubulin signal before photoactivation (Pre-PA), immediately after photoactivation (PA) and at 2 and 4.5 minutes after photoactivation. The PA-GFP-α-tubulin signal was inverted for better visualization. Scale bar is 5 μm. **(B)** Quantitation of kinetochore microtubule (KT-MT) half-life for all cell lines. The first column corresponds to the parental cell line, and the remaining columns correspond to experiments performed after CLASPs depletion with siRNA #12A. Each data point represents an individual cell. The boxes represent median and interquartile interval, the bars represent minimum and maximum values. Parental without RNAi = 43 cells, 5 independent experiments; Parental with RNAi = 20 cells, 6 independent experiments; WT = 67 cells, 7 independent experiments; 2ea-3eeaa = 105 cells, 8 independent experiments; IP12 = 108 cells, 9 independent experiments; IP12-3eeaa = 55 cells, 4 independent experiments; ΔC = 60 cells, 4 independent experiments; ΔC-Gcn4 = 34 cells, 8 independent experiments. Mutant constructs in the TOG domains and in the SxIP motifs, as well the kinetochore-lacking constructs ΔC and ΔC-Gcn4 showed less stable kinetochore microtubules in the absence of endogenous CLASPs. ns = non-significant, * = p<0.05, ** = p< 0.01. **(C)** Quantitation of non-KT-MT half-life for all cell lines for the same data set as in B. All cell lines showed non-KT-MT stability that was indistinguishable from controls. ns = non-significant.

The incapacity to rescue normal kinetochore microtubule half-life after CLASPs depletion was maximal in both C-terminal deletion mutants (Fig. 5A, B). This indicates that the localization of CLASP2 at kinetochores is critical for normal kinetochore microtubule dynamics, independently of CLASP2 dimerization, association with the microtubule lattice and tracking of growing microtubule plus ends. Remarkably, none of the CLASP2 mutants significantly compromised non-kinetochore microtubule half-life in our assay. We conclude that the role of CLASPs in the regulation of spindle length, chromosome congression, error correction and timely SAC satisfaction relies on its capacity to integrate critical microtubule-binding properties at the kinetochore-microtubule interface that sustain microtubule growth and promote their stabilization. These results are consistent with the unique capacity of human CLASPs to bind to curved microtubule protofilaments (Leano et al., 2013; Maki et al., 2015) and recent electron microscopy work showing that both growing and shrinking microtubule plus ends, including those of kinetochore microtubules, are curved (McIntosh et al., 2018). They are also supported by recent in vitro reconstitution experiments that showed that CLASP2-coated spherical beads are able to mediate moderately long-lived attachments with polymerizing microtubule plus-ends (Chakraborty et al., 2019). Unlike other TOG-domain containing proteins, such as members of the XMAP215/ch-TOG family that work as microtubule polymerases that bind to free α,β-tubulin heterodimers (Ayaz et al., 2012; Geyer et al., 2018; Nithianantham et al., 2018), our data is most consistent with a model in which microtubule lattice binding and EB protein-dependent microtubule plus-end tracking of CLASP2 is required at the kinetochore to stabilize, or prevent destabilization, of kinetochore-microtubule attachments as cells progress into metaphase.

## Acknowledgements

We would like to thank Bernardo Orr, Jorge Ferreira and António Pereira for the critical reading of this manuscript. H.G. was supported by a studentship from Fundação para a Ciência e a Tecnologia (SFRH/BD/141066/2018). This work was supported by a Grant-in-Aid for Scientific Research 22570190 (I.A.) and by the European Research Council (grant agreement No 681443) under the European Union’s Horizon 2020 research and innovation program (H.M.).

## Author contributions

H.G. performed and analyzed all the experiments, except Figure 1, under supervision by N.O. and H.M.. A.C.F and Z.G. performed the biochemical characterization of CLASP2, under supervision of S.M.R., J.A. and H.M.. I.A. provided tools. T.M.S. generated tools and preliminary data. H.M. conceived and coordinated the project, designed experiments and analyzed the data. H.G. and H.M. wrote the paper, with contributions from all the authors.

## Material and Methods

### Production of recombinant CLASP2 C-terminal protein

Synthetic gene coding for the C-terminal comprising nucleotides 1213-1515 of human CLASP2α with codon usage optimized for expression in *E. coli* was obtained from GenScript. CLASP2 ORF was subcloned into the *EcoRI* and *NdeI* restriction sites of the expression vector pPR-IBA2 (IBA LifeSciences) in fusion with an N-terminal Strep-tag and a C-terminal His10-tag. *E.coli* BL21 Star (DE3) cells (Life Technologies) transformed with pPR-IBA2-CLASP2 (1213-1515) plasmid were grown at 37 °C in LB medium supplemented with 50 μg/ml ampicillin to OD600 0.4, and expression was induced by addition of 0.4 mM IPTG. After growing for 4 h at 37 °C, cells were harvested and lysed by sonication in 50 mM Tris-HCl pH 8.0, 150 mM NaCl supplemented with protease inhibitors (Complete EDTA-free, Roche). Clarified protein extracts were loaded onto a HisTrap HP column (GE Healthcare) pre-equilibrated in 50 mM Tris-HCl pH 8.0, 500 mM NaCl, 20 mM imidazole (buffer A) and eluted with 200 mM imidazole in buffer A. CLASP2 C-term containing fractions were pooled, adjusted to 100 mM Tris-HCl pH 8.0 and 1 mM EDTA and loaded onto a StrepTrap HP column (GE Healthcare) pre-equilibrated in 100 mM Tris-HCl pH 8.0, 150 mM NaCl, 1 mM EDTA (buffer B). Bound CLASP2 was eluted with 2.5 mM desthiobiotin in buffer B. Protein-containing fractions were concentrated and further purified on a HiPrep 16/60 Sephacryl S-200 HR column (GE Healthcare) pre-equilibrated with 50 mM Tris-HCl pH 8.0, 400 mM NaCl, 5% GlyOH, 1mM EDTA (buffer C). Pure protein was concentrated using a 30 KDa molecular-weight cutoff ultracentrifugal concentration device (Millipore).

### Size exclusion chromatography

Size exclusion chromatography experiments with HeLa cell extracts were performed with a pre-packed Superose 6 10/300 GL column (GE Healthcare ©). The column was equilibrated with buffer A (20 mM Hepes, 150 mM NaCl, 1mM DTT, pH 7.9) and a 0.3 mL/min flow rate was used throughout the experiments. Calibration was performed with protein standards (Stokes radius (a)): Thyroglobulin (8.50 nm), Ferritin (6.10 nm), Aldolase (4.81 nm), BSA (3.55 nm), Ovoalbumin (3.05 nm), Chymotrypsinogen (2.09 nm) and Ribonuclease (1.37 nm). 50 μg of standards and 30 μg of sample cell extract were used in each run. Standard protein elution was monitored by measuring the absorbance at 280 nm. In the sample run, 0.5 mL fractions were collected, TCA precipitated, subjected to SDS-PAGE, blotted and immunodetected with a rat anti-CLASP2 antibody (Maffini et al., 2009), and protein distribution was analyzed by densitometry. Recombinant CLASP2 C-term (300 μg, injection volume of 100 μl) was analyzed on a Superose 12 10/300 GL column (GE Healthcare) pre-equilibrated in buffer C. The column was calibrated with protein standards of known molecular weight and stokes’ radius (Low and High Molecular Weight Kit, GE Healthcare) according to manufacturer’s protocol.

### Density gradient centrifugation

Continuous gradient of 5% and 30% sucrose was prepared in buffer A. 50 μg of each standard protein were loaded on top of the gradient and centrifuged at 29.000 rpm in a SW41 rotor (Beckman) for 29 h at 4°C. After centrifugation, 0.7 mL fractions were collected from the bottom of the tubes, subjected to SDS-PAGE, Coomasie staining and densitometry analysis. Native protein standards used were (Sedimentation coefficient, S) Ribonuclease (2S), Chymotrypsinogen (2.6S), Ovoalbumin (3.5S), albumin (4.6S), immunoglobulins (7S) and catalase (11.3S). 300 μg of sample cell extract were loaded on top of another gradient, and processed as before. After centrifugation, 0.7 mL fractions were collected from the bottom of the tube, TCA precipitated, subjected to SDS-PAGE, blotted and immunodetected with a rat anti-CLASP2 or mouse anti-α-tubulin antibodies (clone B-512, Sigma Aldrich). For analysis of recombinant CLASP2 C-term, density gradients from 5 to 30% sucrose in 50 mM Tris- HCl pH 8.0, 400 mM NaCl, 1mM EDTA were prepared using a gradient former. The gradients were loaded with 50 μg of recombinant CLASP2 C-term or protein standards of known sedimentation coefficient and centrifuged for 29 h at 29.000 rpm in a SW41 rotor (Beckman). Fractions of 700 μl were collected starting from the bottom of the tube using a peristaltic pump. Each fraction was TCA precipitated and analyzed by SDS-PAGE followed by PageBlue Protein Staining (Fermentas).

### Estimation of molecular weight by size exclusion chromatography and density gradient centrifugation

The molecular weights of recombinant CLASP2 C-term and full length CLASP2 were calculated by the method of Siegel and Monty (Siegel and Monty, 1966) using equation 1 and 2 and the respective Stokes radius and sedimentation coefficient obtained from size exclusion chromatography and density gradient centrifugations, respectively,

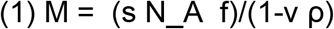

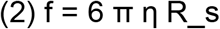

where s is the sedimentation coefficient, NA is the Avogadro’s constant (6.022 x 1023 mol- 1), f is the frictional coefficient, v is the partial specific volume of the protein (0.7362 cm3 g- 1); ρ is the density of the medium (1g cm-3); η is the viscosity of the medium (0.01 g cm-1 s-1) and RS is the stokes radius. The partial specific volume was estimated based on the amino acid sequence of the protein using the method of Cohn and Edsall and the program SEDNTERP v1.08 (http://www.jphilo.mailway.com/default.htm).

### Estimation of molecular weight by dynamic light scattering

Molecular size measurements of CLASP2 C-term (5 mg/ml in buffer C) were performed using a glass cuvette ZEN0023 and a Zetasizer Nano ZS DLS system (Malvern Instruments). Three independent measurements were obtained at 20 °C and data were analyzed using Zetasizer software v7.03. The hydrodynamic radius calculated from the diffusion coefficient using the Stokes-Einstein Equation was determined from the volume distributions and used to estimate the molecular weight.

### Constructs and lentiviral transduction

The mRFP-CLASP2γ constructs WT, IP12 (I496A, P497A; I519A, P520A), 2ea-3eeaa (W106E, K191A; W667E, K833E, K838A, R839A), IP12-3eeaa (I496A, P497A; I519A, P520A; W667E, K833E, K838A, R839A) and ΔC (CLASP2γ without C-Terminal domain 1017-1294), inserted in a CSII-CMV-MCS vector, were reported previously (Maki et al., 2015). The ΔC-Gcn4 construct was generated by inserting the Gcn4 sequence after the mRFP-CLASP2γ-ΔC sequence, using Gibson assembly (Gibson et al., 2009). The entire Gcn4 sequence was contained in the two primers used in the reaction mixed with the vector containing the mRFP-CLASP2γ-ΔC construct cut with *NotI*. Lentiviral supernatants were produced in HEK293T cells using pmd2.G, Pax2 and the CSII-CMV-MLS plasmids, with the presence of the Lipofectamine 2000 (Invitrogen) and Opti-MEM (Gibco). A U2OS cell line stably expressing Photoactivatable (PA) GFP-α-tubulin (Ganem et al., 2005) was transduced with the viral supernatants, in the presence of 6 μg/mL Polybrene (Sigma-Aldrich). The same procedure was used to produce the double expressing H2B-GFP and mRFP-CLASP2γ cell lines.

### Cell line maintenance

All cell lines used in this work were cultured in DMEM with 10% FBS, supplemented with 10 μg/μL of antibiotic-antimycotic mixture (Gibco) and selected with Zeocine 1 μg/μL. Cell lines were maintained at 37°C in a 5% CO_2_, humidified atmosphere. To synchronize HeLa cell cultures in mitosis 5 μM of S-Trytil-L-Cysteine were added to the medium 16 h before harvesting by mitotic shake off.

### Cell sorting

Cells expressing the different mRFP-CLASP2γ constructs were enriched by cell sorting using a FACS Aria II cell sorter (Becton Dickinson). Cells were re-suspended in Basic Sorting Buffer [Phosphate Buffered Saline (PBS) 1X, Ca^2+^ and Mg^2+^ free; 5 mM EDTA; 25 mM HEPES; 2% FBS] and cells expressing red fluorescence signal were sorted to a tube containing DMEM with 10% FBS and then transferred to an appropriate growth flask. The same procedure was used to enrich the double expressing H2B-GFP and mRFP-CLASP2γ cell lines, selecting in cell with double fluorescence signal (GFP and mRFP).

### siRNA

The siRNA experiments were performed in 0.2×10^6^ cells cultured in 1.5 mL DMEM with 5% FBS. A solution containing 2 μL of Lipofectamine RNAiMax (Invitrogen) diluted in 250 μL Opti-MEM was mixed with other containing 100 pmol of siRNA (Sigma Aldrich) diluted in 250 μL Opti-MEM. The solution mix was incubated for 30 minutes at room temperature and then added dropwise to the cells. The siRNA oligonucleotides used were: Scrambled siRNA CUUCCUCUCUUUCUCUCCCUUGUGA, CLASP1#A siRNA GCCAUUAUGCCAACUAUCU, CLASP2#A siRNA GUUCAGAAAGCCCUUGAUG, CLASP1#B siRNA GGAUGAUUUACAAGACUGG and CLASP2#B siRNA GACAUACAUGGGUCUUAGA (Mimori-Kiyosue et al., 2005). After 6 h incubation at 37°C, the medium was removed and replaced by 2 mL of DMEM with 10% FBS.

### Western blotting

Protein extracts were obtained from cells by addition of Lysis Buffer (20 mM HEPES/KOH, 1 mM EDTA, 1 mM EGTA, 150 mM NaCl, 0.5% NP-40, 10% glycerol, 2 mM DTT at −20°C and Protease inhibitor 4C+PMSF 0.1 mM at −20°C 1:100, pH 7.9) and frozen with liquid Nitrogen (N2). The suspension was centrifuged 5 minutes at 14,000 rpm at 4°C, collecting the supernatant. Total protein levels were quantified using Bradford Reagent (Thermoscientific), using Bovine Serum Albumin (BSA; Thermofisher) solutions as standards. 15 μg of total protein per sample were mixed with Sample Buffer (50 mM Tris-HCl pH 6.8, 2% SDS, 10% glycerol, 1% β-mercaptoethanol, 12.5 mM EDTA, 0.02% bromophenol blue) and denatured at 95°C for 5 minutes. Samples were loaded in a 6.5% Acrylamide Gel mounted in a Mini-PROTEAN vertical electrophoresis apparatus (Bio-Rad), using NZYColour Protein Marker II (NZYTech). Blotting was performed with an iBlot Gel Transfer System (Invitrogen). Membranes were blocked with 5% powder milk in PBS Tween 0.1% for 45 minutes. For primary antibodies we used hybridoma supernatant of monoclonal antibody 6E3 Rat anti-CLASP2 (Maffini et al., 2009) at 1:150 and Mouse anti-α-tubulin clone B-512 (Sigma Aldrich) at 1:10000, diluted in 5% powder milk in PBS Tween 0.1%, and incubated overnight (4°C) with agitation. Anti-Rat HRP (Horseradish Peroxidase) and anti-Mouse HRP secondary antibodies (Santa Cruz Biotechnology) were used at 1:2000 proportions in 5% powder milk in PBS Tween 0.1% and incubated for 1 h. Signal was developed with Clarity Western ECL Blotting Substrate (Bio-Rad) and detected in a Bio-Rad Chemidoc XRS system, with Image Lab software.

### Immunofluorescence

Cells were fixed using paraformaldehyde at 4% in Cytoskeleton Buffer (CB; 137 mM NaCl, 5mM KCl, 1.1 mM Na_2_HPO_4_, 0.4 mM KH_2_PO_4_, 2mM EGTA, 2mM MgCl_2_, 5 mM PIPES, 5 mM Glucose), or in alternative, Methanol at −20°C, for 10 minutes (for Nocodazole arrested cell experiments, Nocodazole was added at final concentration of 3.3 μM to the culture, 2-3 h prior to fixation). Extraction was accomplished using CB-Triton X-100 0.5% for 10 minutes. Primary antibodies used in this work were: mouse anti-α-tubulin clone B-512 at 1:1500 (Sigma Aldrich), mouse anti-EB1 at 1:500 (BD Biosciences), human anti-CREST 1:1000 (Abyntek), rat anti-RFP (ChromoTek) 1:1000, diluted in PBS-Triton X-100 0.1% with 10% FBS. Secondary antibodies Alexa Fluor anti-Mouse 488, anti-rat 568 and anti-Human 647 (Invitrogen) were diluted 1:1000 in PBS-Triton X-100 0.1% with 10% FBS. DNA staining was achieved by addition of 1 μg/ml 4',6-diamidino-2-phenylindole (DAPI; Sigma-Aldrich). Coverslips were mounted using mounting medium (20 mM Tris pH8, 0.5 mM N-propyl gallate, 90% glycerol). Imaging was performed in a Zeiss Axio Observer Widefield Microscope, using a 63x 1.46 N.A. plan-apochromatic objective. 3D-deconvolution was performed using AutoQuant X Image Deconvolution Software (Media Cybernetics). Image processing and quantifications were made using ImageJ Software. Statistical analysis was performed using GraphPad Prism 8 Software, using Student t-test or Mann-Whitney test, according to the normality of the distribution.

### Live-cell imaging

The mRFP-CLASP2γ U2OS cells expressing H2B-GFP were cultured in glass coverslips using DMEM without phenol red and supplemented with 25 mM of 4-(2-hydroxyethyl)-1-piperazineethanesulfonic acid (HEPES; Gibco) and 10% FBS. SiR-Tubulin (Spirochrome) was added to a final concentration of 20 nM, 30-60 minutes prior to observation. Time-lapse imaging was performed in a heated chamber (37°C) using a 60x oil-immersion 1.4 N.A. plan-apochromatic objective mounted on an inverted microscope (Nikon TE2000U) equipped with a CSU-X1 spinning-disk confocal head (Yokogawa Corporation of America) and with three laser lines (488 nm, 561 nm and 647 nm). Images were detected with an iXonEM+ EM-CCD camera (Andor Technology). Eleven z-planes separated by 1 μm were collected every 2 minutes. Image processing and quantifications was performed using ImageJ software and NIS Viewer from Nikon. Statistical analysis was performed using GraphPad Prism 8 Software, using Student t-test or Mann-Whitney test, according to the normality of the distribution.

### Photoactivation experiments

Microtubule turnover rates were measured in the mRFP-CLASP2γ U2OS cells stably expressing the PA-GFP-α-tubulin. Cells were grown in glass coverslips with DMEM without phenol red and with 25 mM of HEPES (Gibco), supplemented with 10% FBS. Time-lapse imaging was performed in a heated chamber (37°C) using a 100x 1.4 N.A. plan-apochromatic differential interference contrast (DIC) objective mounted on an inverted microscope (Nikon TE2000U) equipped with a CSU-X1 spinning-disk confocal head (Yokogawa Corporation of America) and with two laser lines (488 nm and 561 nm). Images were detected with an iXonEM+ EM-CCD camera (Andor Technology). Photoactivation was performed with a Mosaic DMD-based patterning system (Andor) equipped with a 405 nm diode laser. Cells in metaphase were chosen using DIC acquisition. A thin stripe of spindle microtubules was locally photoactivated in one half-spindle by pulsed near-UV-irradiation (405 nm laser; 500 microseconds exposure time) and fluorescence images (seven 1 μm-separated z-planes centered at the middle of the mitotic spindle) were captured every 15 seconds for 4.5 minutes with a 100x oil-immersion 1.4 N.A. plan-apochromatic objective. To quantify fluorescence dissipation after photoactivation (FDAPA), spindle poles were aligned horizontally and whole-spindle, sum-projected kymographs were generated (sum projections using ImageJ and kymographs generated as previously described (Pereira and Maiato, 2010). Fluorescence intensities were quantified for each time point (custom-written routine in Matlab, “LAPSO” software) and normalized to the first time point after photoactivation for each cell following background subtraction and correction for photobleaching. Correction for photobleaching was performed by normalizing to the values of fluorescence loss obtained from whole-cell (including the cytoplasm), sum projected images for each individual cell. This method allows for the precise measurement of photobleaching for each cell at the individual level and avoids any potential unspecific Taxol-associated phenotypes. Under these conditions, the photoactivated region dissipates, yet the photoactivated molecules are retained within the cellular boundaries defined by the cytoplasm. To calculate microtubule turnover, the sum intensity at each time point was fit to a double exponential curve A1*exp(−k_1_*t) + A2*exp(−k_2_*t) using Matlab (Mathworks), in which *t* is time, *A1* represents the less stable (non-KT-MT) population and *A2* the more stable (KT-MT) population with decay rates of *k*_1_ and *k_2_*, respectively. When running the routine to fit the data points to the model curve, the rate constants are obtained as well as the percentage of microtubules for the fast (typically interpreted as the fraction corresponding non-kinetochore microtubules) and the slow (typically interpreted as the fraction corresponding to kinetochore microtubules) process are obtained. The half-life for each process was calculated as ln2/*k* for each population of microtubules. All experiments were performed in the presence of MG132 (5 μM) added 30 min prior to imaging to ensure that cells were in metaphase and prevent mitotic exit. Statistical analysis for photoactivation experiments was performed in the GraphPad Prism8 software, using the Mann-Whitney test.

### Statistical Analysis

Statistical analysis was performed with Graphpad Prism, version 8. Quantification of kinetochore co-localization, spindle length, segregation errors, NEB to Anaphase and NEB to Metaphase durations were performed using the parametric unpaired Student’s t-test. Quantification of both kinetochore microtubule and non-kinetochore microtubule half-lives was performed using the non-parametric Mann-Whitney U-Test.

---

**Supplementary Figure 1.** Western blot with anti-CLASP2 antibody, revealing the total (endogenous and exogenous) CLASP2 levels for each mRFP-CLASP2γ construct cell line. Endogenous CLASP2α and the mRFP-CLASP2γ constructs WT, 2ea-3eeaa, IP12 and IP12-3eeaa could not be distinguished due to similar molecular weight (165.7 kDa for endogenous CLASP2α and 167.9 kDa for mRFP-CLASP2γ constructs), while in the ΔC (120.8 KDa) and ΔC-Gcn4 (125.3 KDa) it was possible to visualize both bands for endogenous CLASP2α and exogenous mRFP-CLASP2γ. All cell lines expressed comparable levels of mRFP-CLASP2γ, slightly lower in 2ea-3eeaa and slightly higher in ΔC. α-tubulin was used as loading control.

**Supplementary Figure 2.** Western blot analysis, using an anti-CLASP2 antibody, highlighting the depletion of endogenous CLASPs by siRNA #12A and siRNA #12B. Since endogenous CLASP2α and the mRFP-CLASP2γ constructs WT, IP12, 2ea-3eeaa and IP12-3eeaa displayed similar molecular weights (MW) (165.7 kDa for endogenous CLASP2α and 167.9 kDa for the mRFP-CLASP2γ constructs), depletion efficiency of endogenous CLASPs were obtained by comparing the treatments with Scramble siRNA, siRNA #12A that deplete endogenous CLASPs and siRNA #12B that deplete both endogenous and exogenous CLASPs. However, for ΔC and ΔC-Gcn4 cases, because they have different MW from endogenous CLASP2α, it was possible to discriminate both bands. α-tubulin was used as loading control.

**Supplementary Figure 3. (A)** Immunofluorescence analysis showing the co-localization of the mRFP-CLASP2γ constructs with EB1 at the microtubule plus ends in interphase. Cells were stained with anti-EB1 antibody (green) and anti-RFP antibody (red). The anti-RFP antibody was used in order to increase detection of the mRFP-CLASP2γ constructs at the plus tips after methanol fixation. IP12 and IP12-3eeaa showed a decreased co-localization with EB1 at the microtubule plus ends. Scale bar is 5 μm. **(B)** Immunofluorescence analysis showing localization of the mRFP-CLASP2γ constructs with mitotic spindle microtubules. Cells were immunostained with an anti-α-tubulin antibody to visualize the spindle (green). The 2ea-3eeaa and IP12-3eeaa constructs showed a decreased localization on spindle microtubules. Scale bar is 5 μm.

## Movie Legends

**Movie S1 –** Human parental U2OS cell stably expressing PA-GFP-α-tubulin (not shown) and H2B-GFP (green in the merged image; shown alone in white in the right panel) after CLASP1 and CLASP2 RNAi, with microtubules (magenta) visualized with 20 nM SiR-tubulin. Note the presence of a lagging chromosome during anaphase. Time is in h:min.

**Movie S2** – Human parental U2OS cell stably expressing PA-GFP-α-tubulin and wild type (WT) mRFP-CLASP2γ (red) and H2B-GFP (green in the merged image; shown alone in white in the right panel) after CLASP1 and CLASP2 RNAi, with microtubules (magenta) visualized with 20 nM SiR-tubulin. Time is in h:min.

**Movie S3** – Human parental U2OS cell stably expressing PA-GFP-α-tubulin and mRFP-CLASP2γ 2ea-3eeaa mutant (red) and H2B-GFP (green in the merged image; shown alone in white in the right panel) after CLASP1 and CLASP2 RNAi, with microtubules (magenta) visualized with 20 nM SiR-tubulin. Note the presence of 2 lagging chromosomes during anaphase, one of which resolves. Time is in h:min.

**Movie S4** – Human parental U2OS cell stably expressing PA-GFP-α-tubulin and mRFP-CLASP2γ IP12 mutant (red) and H2B-GFP (green in the merged image; shown alone in white in the right panel) after CLASP1 and CLASP2 RNAi, with microtubules (magenta) visualized with 20 nM SiR-tubulin. Note the presence of a chromosome bridge in anaphase and telophase. Time is in h:min.

**Movie S5** – Human parental U2OS cell stably expressing PA-GFP-α-tubulin and mRFP-CLASP2γ IP12-3eeaa mutant (red) and H2B-GFP (green in the merged image; shown alone in white in the right panel) after CLASP1 and CLASP2 RNAi, with microtubules (magenta) visualized with 20 nM SiR-tubulin. Note the presence of a transient lagging chromosome during anaphase. Time is in h:min.

**Movie S6** – Human parental U2OS cell stably expressing PA-GFP-α-tubulin and mRFP-CLASP2γ ΔC mutant (red) and H2B-GFP (green in the merged image; shown alone in white in the right panel) after CLASP1 and CLASP2 RNAi, with microtubules (magenta) visualized with 20 nM SiR-tubulin. Note the presence of transient lagging chromosomes in the beginning of anaphase. Time is in h:min.

**Movie S7** – Human parental U2OS cell stably expressing PA-GFP-α-tubulin and mRFP-CLASP2γ ΔC-Gcn4 mutant (red) and H2B-GFP (green in the merged image; shown alone in white in the right panel) after CLASP1 and CLASP2 RNAi, with microtubules (magenta) visualized with 20 nM SiR-tubulin. Note the presence of persistent lagging chromosomes in anaphase. Time is in h:min.

