## Supplementary figures and images for "CLASP2 lattice-binding near microtubule plus ends stabilizes kinetochore attachments"

### Figure S1

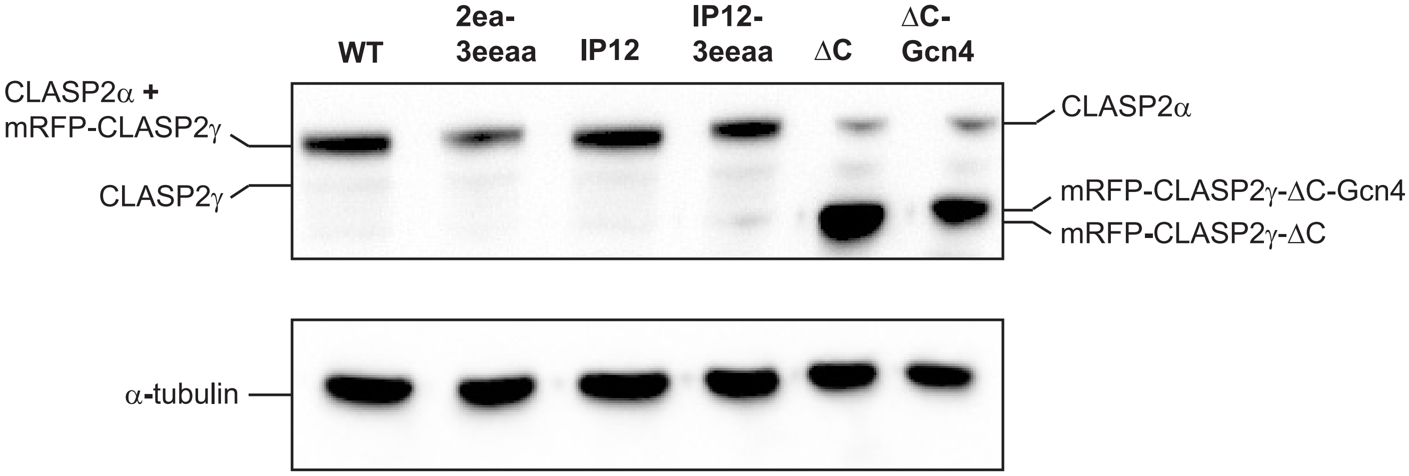

### Figure S2

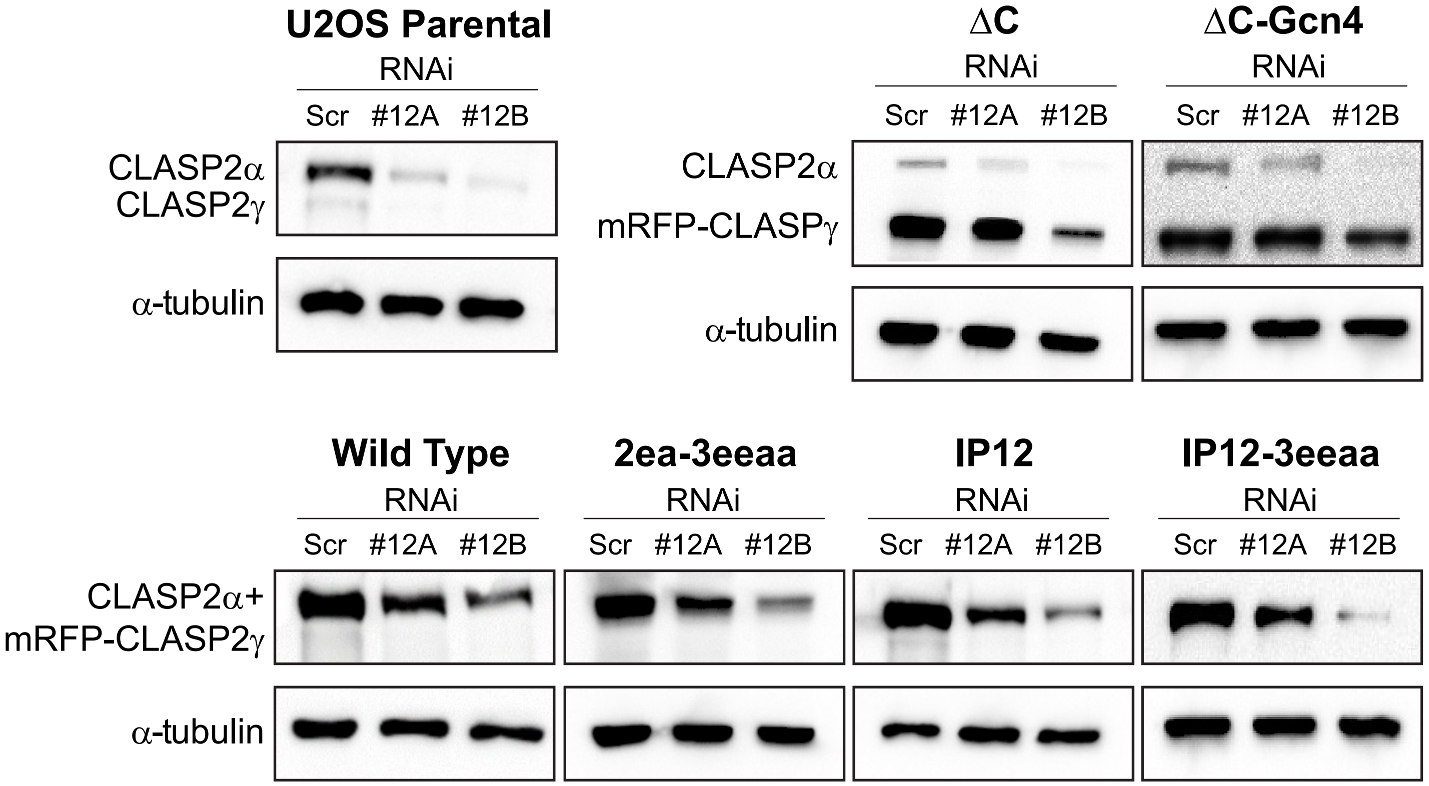

### Figure S3

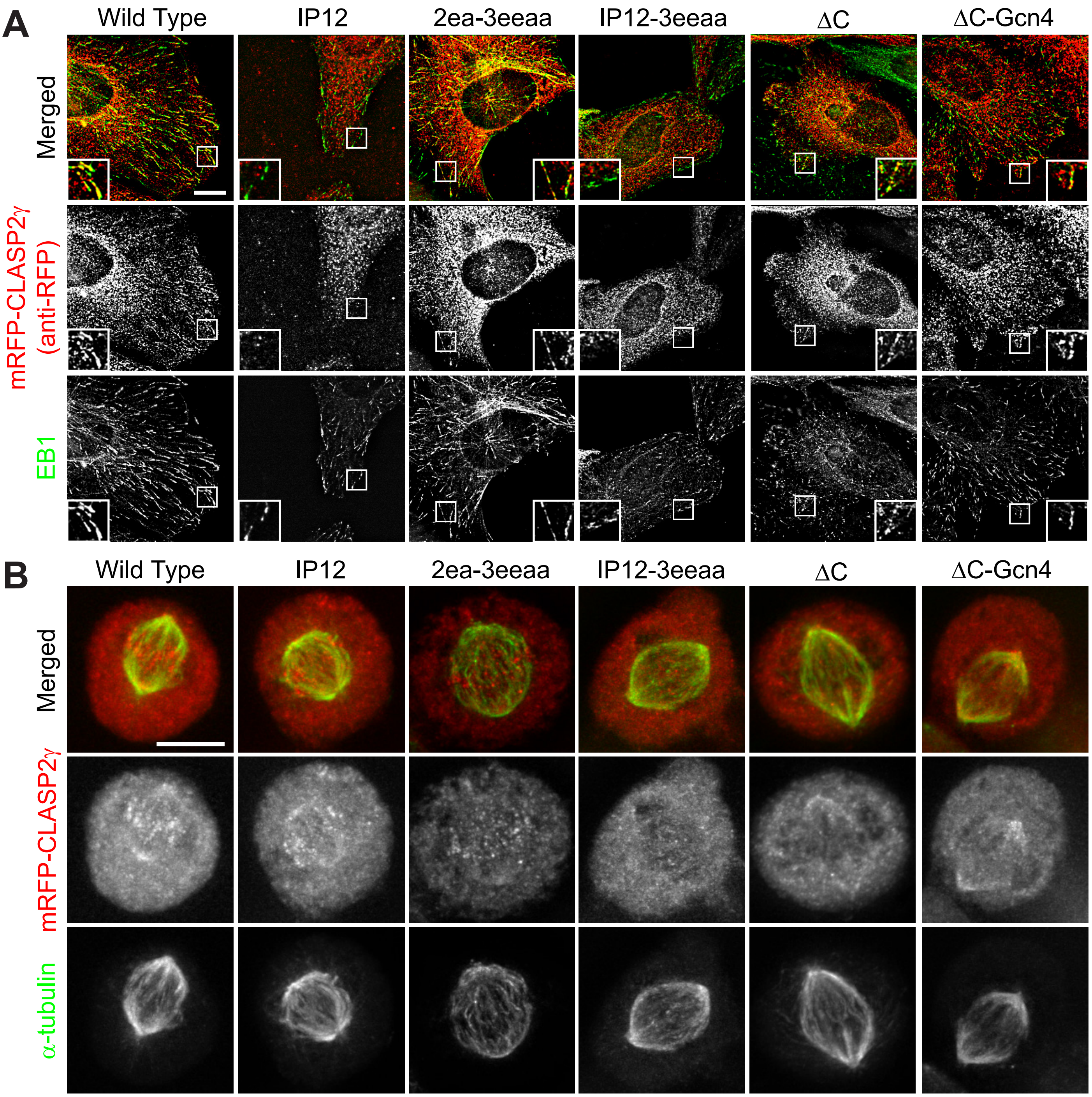
